# ODOR IDENTITY CAN BE EXTRACTED FROM THE RECIPROCAL CONNECTIVITY BETWEEN OLFACTORY BULB AND PIRIFORM CORTEX IN HUMANS

**DOI:** 10.1101/2021.03.18.436041

**Authors:** Behzad Iravani, Artin Arshamian, Mikael Lundqvist, Leslie Kay, Donald A. Wilson, Johan N. Lundström

## Abstract

Neuronal oscillations route external and internal information across brain regions. In the olfactory system, the two central nodes—the olfactory bulb (OB) and the piriform cortex (PC)—communicate with each other via neural oscillations to shape the olfactory percept. Communication between these nodes have been well characterized in non-human animals but less is known about their role in the human olfactory system. Using a recently developed and validated EEG-based method to extract signals from the OB and PC sources, we show in healthy human participants that there is a bottom-up information flow from the OB to the PC in the beta and gamma frequency bands, while top-down information from the PC to the OB is facilitated by delta and theta oscillations. Importantly, we demonstrate that there was enough information to decipher odor identity above chance from the low gamma in the OB-PC oscillatory circuit as early as 100ms after odor onset. These data further our understanding of the critical role of bidirectional information flow in human sensory systems to produce perception. However, future studies are needed to determine what specific odor information is extracted and communicated in the information exchange.

## INTRODUCTION

Communication within and between neural circuits is facilitated by oscillations in neural activity across a broad spectrum of frequencies (Bonnefond et al. 2017; Buzsáki 2006; Fries 2005; Fries 2015; Hipp et al. 2011; Varela et al. 2001). In human and animal models alike, this oscillatory activity has been shown to support sensory coding, memory, and attention (Lakatos et al. 2008). The mammalian olfactory system was one of the earliest systems where such oscillatory activity was described, specifically within the olfactory bulb (OB) (Adrian 1942; Adrian 1950; Freeman 1959; Freeman 1972; Freeman 1974). Subsequent studies in model species have demonstrated a role for oscillations within the whole olfactory pathway [e.g., the piriform cortex (PC)] and related structures [e.g., hippocampus] (Kay 2014; Wilson and Sullivan 2011). One fundamental role that neural oscillations serve is entrainment of activity across different regions which amplifies information flow (Bonnefond et al. 2017; Buzsáki 2006; Fries 2005; Fries 2015; Hipp et al. 2011; Varela et al. 2001). This entrainment is especially important for olfactory processing were the information flow between connected regions, such as the OB and PC, is reciprocal with beta being bottom-up connection but with overturned directionality during odor sampling (Gourévitch et al. 2010; Kay and Beshel 2010). Particularly, top-down signal flow conveys information about expectation, state, memory, or attention, which, in turn, shape beta oscillations, and more comprehensively the stimulus encoding in the more peripheral OB (Gourévitch et al. 2010; Kay and Beshel 2010; Martin and Ravel 2014; Wilson and Yan 2010). Similar events occurs in thalamocortical sensory systems, wherein the sensory cortex can modulate thalamic sensory-driven output based on context and task demands (Guillery and Sherman 2002). Although well studied in rodent models, significantly less is known about the role of odor-evoked oscillations in the early processing stages of the human olfactory system.

In one of the few studies on the oscillatory activity in human olfaction, Jiang and colleagues (2017) recently demonstrated not only that oscillatory activity in the theta band conveyed information odor within 500 milliseconds of the sniff onset in the PC, but also demonstrated an increased theta-specific phase coupling between the PC and hippocampus during the same temporal window. This clearly demonstrates that, akin to what has previously been demonstrated in animal models, oscillation-dependent communication is an important communication tool also within the early human olfactory system. To date, no study has assessed oscillatory communication between the OB and PC in human participants which constitutes a significant gap in our knowledge of the olfactory system given that the OB has been suggested to fulfil a role comparable to both V1 (Shepherd et al. 2004) and the thalamus (Kay and Sherman 2007) in the visual system. This lack of data can be attributed to the fact that it has not been technically possible to obtain non-invasive and temporally precise recordings from the human OB until a recent EEG-based methodological development — the electrobulbogram (EBG) — that enables a direct and non-invasive functional measure of the human OB (Iravani et al. 2020). This method, that was recently validated in a range of experiments as a reliable method to assess signal from the human OB (Iravani et al. 2020), has demonstrated that, like its non-human animal counterpart, the human OB generates gamma oscillations during odor processing (Iravani et al. 2020).

Here, using this validated and non-invasive recording technique that allows us to acquire odor-induced activity within both the human OB and PC during passive odor perception, we seek to answer the fundamental question whether reciprocal oscillatory connectivity between the OB and the PC also exists in humans. We identify a unique oscillatory bottom-up and top-down information flow in the OB-PC circuit. Importantly, we demonstrate that this OB-PC communication predicts odor identity^1^.

## METHOD

### Participants

A total of 29 healthy individuals (mean age = 27 ± SD 5.30, 18 women) participated in the study. All participants were non-smokers and had no history of head trauma leading to unconsciousness, nor any reported past neurological problems. A functional sense of smell was assessed using a 5-items 4-alternative cued odor identification task prior to EEG recording with all participants passing the criteria for inclusion with at least 3 correct answers. Considering the age range of our sample and given our inclusion criteria, the chance of erroneously including individuals with functional anosmia in the experiment was less than 0.05% with this screening method. Participants avoided eating and drinking anything other than water for at least 6 hours before testing to maximize electrophysiological response from the olfactory bulb (Iravani et al. 2020). The study was approved by the Swedish Ethical Review Authority and signed informed consent from participants was obtained prior to participation.

### Chemicals and odor delivery

To increase the ecological validity of the stimuli, we used two odor mixtures of food odors and one monomolecular non-food odor. These were Orange (Sigma Aldrich, # W282510, CAS 8008-57-9), Chocolate (Givaudan, VE00185273), and n-Butanol (Merck, CAS 71-36-3) diluted to 30%, 15%, and 20%, respectively, in neat diethyl phthalate (99.5% pure, Sigma Aldrich, CAS 84-66-2). The dilution values are given as volume/volume from neat compounds and concentration for each odor was determined in a pilot experiment with the aim of producing iso-intense odors of equal pleasantness and familiarity. Participants in this experiment rated each odor and, as evident in **Fig 1**, average odor intensity (BF_01_ = 0.34) and familiarity (BF_01_ = 0.25) were largely similar across odors within all three perceptual categories determined by Bayesian repeated measured ANOVA. However, there was strong evidence for rejecting the null hypothesis (BF_01_ = 0.04) for pleasantness, indicating that odors were significantly different in pleasantness. In the Bayesian analysis, the prior for the fixed effect was a normal distribution with mean of 0 and standard deviation 0.2. For the random effect a half Cauchy with scaling factor of 0.5 was considered as the prior.

**Fig 1.**
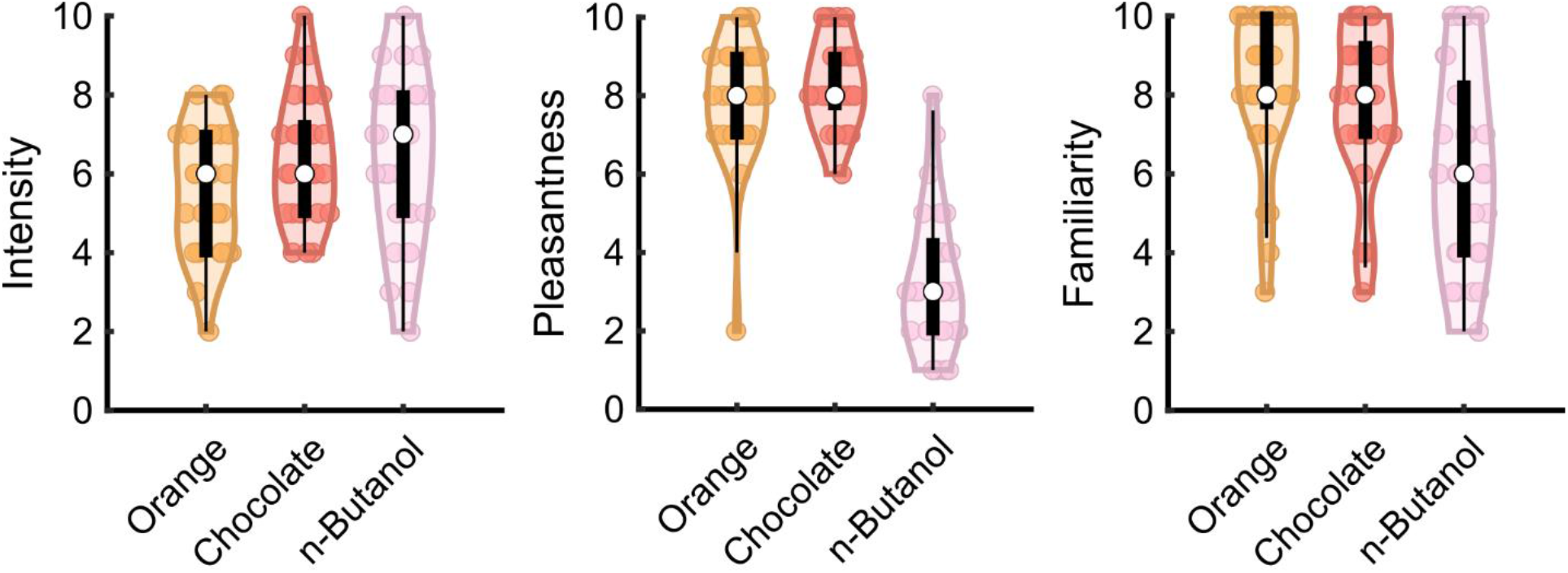
Odor perceptual ratings. Intensity, pleasantness, and familiarity scores of odor stimuli. Violin plots show the distribution of the ratings. The white dot shows the median and the black box shows %75 and %25 quantiles. Individual data points are marked with circles in respective color. The whiskers show the extreme value of the data points reaching 1.5 times the inter quartile range above 75% or below 25% quantiles.

All odors were birhinally delivered in a random order with a total flow rate of 3L/min for a length of 1s (condition: Odor) using a short rise-time (i.e. less than 200ms) computer-controlled olfactometer (Lundström et al. 2010). Interspersed with the odor trials, there were trials consisting of 1s long 3L/min clean air stimuli to assess both neural processing during no odor nasal inhalation (condition: Clean Air) and potential tactile sensations caused by air fluctuation at the onset of a trial due to valve switching. After each trial, participants performed a four-alternative force-choice task containing the name of the correct odor choice, two distractor odor object names, and the option to select ‘no odor’. The odor identification answers from one individual were excluded due to a corrupt data file.

Moreover, to limit potential unintended tactile stimulation at the onset of a stimulus, a constant clean air flow of 0.3 L/min was maintained during the whole experiment and stimuli were inserted into the ongoing flow. Hence, considering the constant flow and conditions’ flow rate, a total airflow of 3.3 L/min was held constant during the trials which yield 1.65 L/min per nostril after the flow is delivered to the individual nostril, a flow well below the threshold known to cause nasal irritation (Lundström et al. 2010). Additionally, we prevented potential effects of onset expectation by implementing a nasal inhalation-triggered design in which all trials’ onset were synchronized in phase with inhalation and unbeknown to the participant. Given that the activity of 50% of all Mitral and tufted cells in the rodent OB are intertwined with respiration (Kay and Laurent 1999), aligning onset of trials to onset of the inhalation further increased our signal-to-noise ratio (SNR). We enabled inhalation triggering by tracing the inhalation pattern using a small nasal temperature probe attached next to the right nostril (MLT415, ADInstrument, Colorado). A trigger-threshold, individually set for each participant, triggered the olfactometer slightly before the nadir of the respiratory cycle and consequently matched stimuli onset (factoring in the known rise-time) with nasal inspiration. Temperature change was sampled at the rate of 400 Hz (Powerlab 16/35, ADInstruments, Colorado) and processed in LabChart Pro version v7.3.8. Subsequently we assessed the length and the area under the curve of inhalations across conditions where we found no difference, determined by repeated measures ANOVA, for neither length F(3,84) = 0.74, p > .53 nor area under the curve of inhalation F(3,84) = 1.00, p > .39 (supplementary **Fig S1**).

Timing and stimulus triggering were implemented within E-prime 2 (Psychology Software Tools, Pennsylvania). All the recordings were carried out in a sound attenuating and shielded booth for psychophysical testing with high air turnover rate to vent out potential lingering odors. Participants wore headphones with low-level white noise played through them during the whole experiment to avoid potential unintended auditory onset cues due to air flow from the olfactometer. The volume of the noise was individually adjusted to maintain participants’ comfort during the test. A jittered pre-stimulus interval (600 – 2000 ms) was added before the onset of each trial to further minimize predictability of odor onset by participants. Moreover, to limit odor habituation effects, a long average inter-trial interval (14000 ms) was used.

### Electroencephalography, Electrobulbogram, neuronavigation data collection

We collected data from 64 scalp EEG electrodes together with 4 electrobulbogram (EBG) electrodes above the eyebrows (Iravani et al. 2020). Signals were sampled at 512 Hz using an active-electrode EEG system (ActiveTwo, BioSemi, Amsterdam, The Netherlands). Prior to recording, we visually controlled the electrodes’ offset and adjusted those above 40mV until the offset met the satisfactory threshold value. EEG scalp electrode placement followed the international 10/20 standard and later during analysis were re-referenced to average of all electrodes.

Following the attachment of all electrodes, we digitized their position in stereotactic space using an optic neuro-navigation system (BrainSight, Rogue Research, Montreal, Canada). We implemented the digitization of electrodes’ position by localizing fiducial landmarks such as the nasion and left/right preauricular as well as the central point of each electrode. We used these landmarks to co-register electrode coordinates to the standard MNI space. The digitized electrode positions were later used in the eLORETA algorithm to project sensor data into source space.

### Experimental design and Statistical Analyses

EEG/EBG signals pre-processing was started by epoching the data from 500 ms pre-stimulus to 1500 ms post-stimulus, followed by re-referencing to the average of all electrodes, band-pass filtering at 1-100 Hz and power line-filtering at electrical frequency using DFT filters (Iravani et al. 2020). The re-referencing to averaged electrodes enabled us to estimate an un-biased source activity. In total, 135 trials were recorded from each individual among which there were 35 trials for each odor and 30 trials for Clean Air. Additionally, we detected trials with muscle and eye-blink artifacts, using an automatic artifact rejection algorithm. In brief, for implementing the algorithm, we band-passed the data at frequency ranges susceptible to each specific artifact and estimated the amplitude using Hilbert transform followed by Z-scoring. We removed trials with Z-values more than 6 for muscle, and 4 for blink, artifacts at susceptible frequencies [for more details, please see (Iravani et al. 2020)]. Finally, a visual inspection was carried out to remove trials with exceedingly high variance.

### Source reconstruction time-course

Source reconstruction, given a head model and a source model, allowed us to extract OB and PC time-courses. We reconstructed source space time-courses using eLORETA algorithm with common spatial filter approach, thus we used a common solution to reconstruct Odor and Clean Air, time-series in source space. The covariance between electrodes for 1 second of stimulus presentation (i.e., Odor and Clean Air) was calculated. Then, we constructed a spherical head-model with four spheres (i.e., scalp, skull, grey matter and white matter) based on the MNI template with conductivity of 0.43, 0.01, 0.33, and 0.14 (**Fig 2**). A grid with 1cm spacing was used for source modelling. We placed dipoles on the grid points where grey matter probability was larger than 40%. Digitized electrode positions were co-registered to the head-model using a six-parameter affine transformation. Next, the sensor time-courses were transformed into the source space as a cross production of spatial filter, estimated by eLORETA, and the sensor time-course. Later, we constrained our analysis into two ROIs where the dipoles corresponded to the left OB (x −6, y 30, z −32), right OB (x 6, y 30, z −32) (Iravani et al. 2020), left PC (x −22, y 0, z −14) and right PC (x 22, y 2, z −12) (Seubert et al. 2013). Trials with muscle and blink artifacts were subsequently removed and time-courses across hemispheres were averaged (**Fig 2**). Finally, the source activity was projected to the principal axis using singular value decomposition. Notably, the regularization parameter was set to 10% and applied before decomposition of covariance matrix in eLORETA. The source reconstruction was performed using field trip toolbox 2018 within Matlab R2019b (Oostenveld et al. 2011).

**Fig 2.**
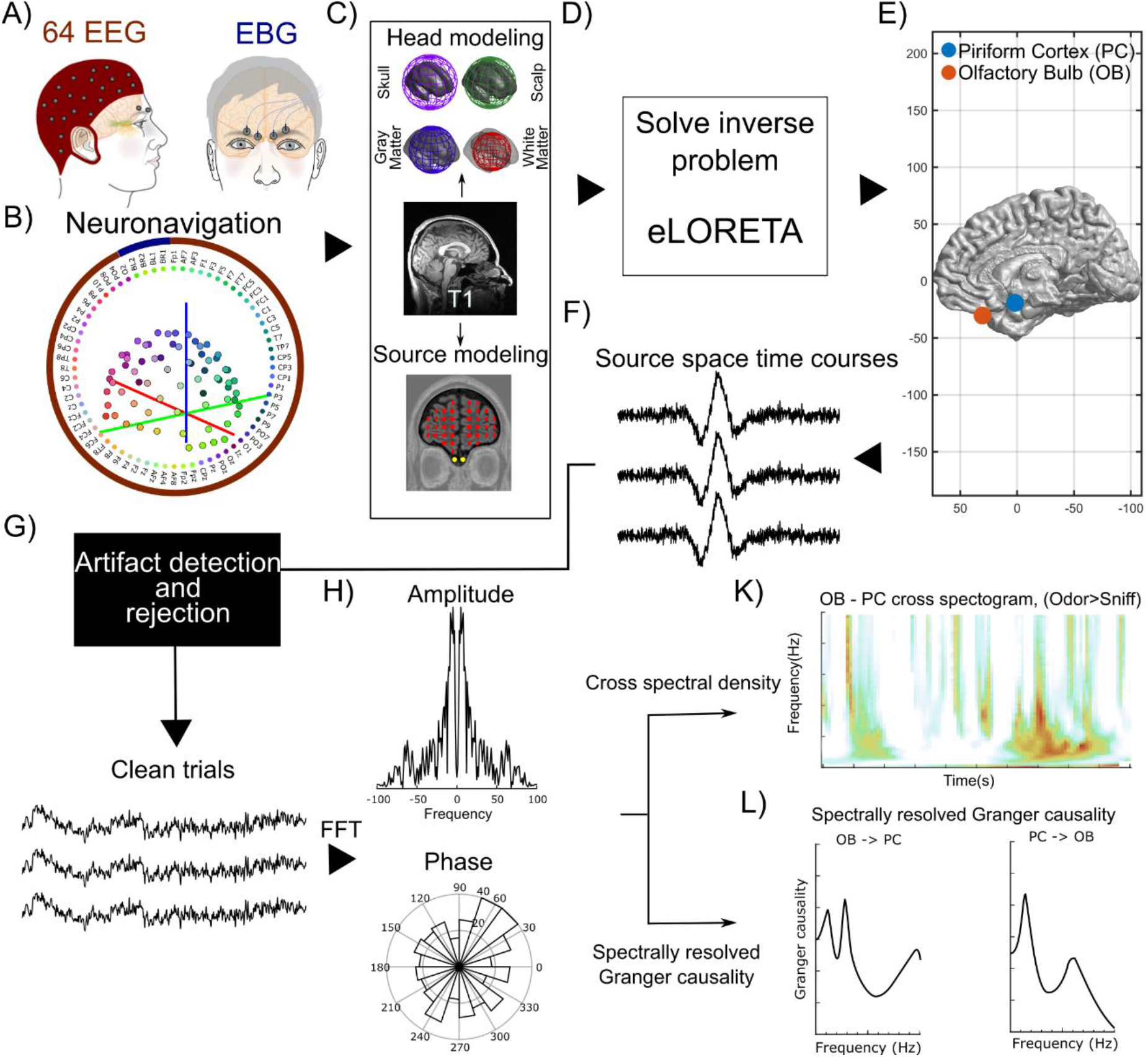
Summary of the analysis procedure. (**A**) Data from 64 EEG together with 4 EBG electrodes were collected from individuals during an odor identification task. (**B**) Electrode positions were digitalized using an optical neuronavigation system to be later used for source reconstruction. (**C**) Using MNI T1 MRI template, a spherical head model, including 4 concentric spheres representing different head tissue, and underlying source model was created. (**D**) Neuro navigated electrode position, head model, and source model were fed into the eLORETA algorithm to reconstruct source time-courses. (**E**) Dipoles corresponding to olfactory bulb (OB) and piriform cortex (PC) were identified. (**F**) Time-course activity of OB and PC were extracted. (**G**) Trials with artifacts were identified and removed from further analysis. (**H**) Clean trials of OB and PC time-courses were transformed in Fourier domain. FFT denotes fast Fourier transform. (**K**) The OB-PC connectivity was quantified as cross spectrogram. (**L**) The effective connectivity of OB-PC was assess using spectrally resolved Granger causality.

Extracting signal from estimated source location is always susceptible to signal loss. To assess the reconstructed signal quality, the mean amplitude of the source time course was estimated by applying a Hilbert transform followed by averaging of envelope signal over 1s stimulus interval and converting to decibels (dB). The difference between the mean amplitude of Odor and Clean Air trials were used as estimation of odor SNR. SNRs were subsequently sorted by their physical depth from the cortical surface and the 95% confidence interval (CI) of SNR at the depth corresponding to PC (i.e. 80 to 100mm) was calculated. Finally, we assessed the SNR at PC by comparing with 0, i.e. where the mean amplitude of Odor is equal to Clean Air.

### PC reconstructed time-course odor SNR is significantly above noise level

In previous work, we found the OB activity can be reliably measured using EBG electrodes, but it is of interest to also assess the validity and quality of reconstructed PC time-course. Using sensitivity analysis, we assessed the odor signal-to-noise ratio (SNR) of potential sources at various depths and compared the mean amplitude of Odor trials versus Clean Air trials as a function of source depth. A 3D grid with 1cm spatial resolution was co-registered to the default MNI brain template where a potential source was considered for the grid’s vertex if the gray matter probability of that point was above 40%. We found that the odor SNR was well above the noise level for both the left PC, *t*(28) = 8.53, *p* < 3e-9, *CI* = [0.21 0.34], and the right PC, *t*(28) = 7.44 *p* < 4e-8, *CI* = [0.18 0.30], given the 95% confidence interval of noise at the depth corresponding to PC, as well as situated in close proximity within the archived space (**Fig 3**).

**Fig 3.**
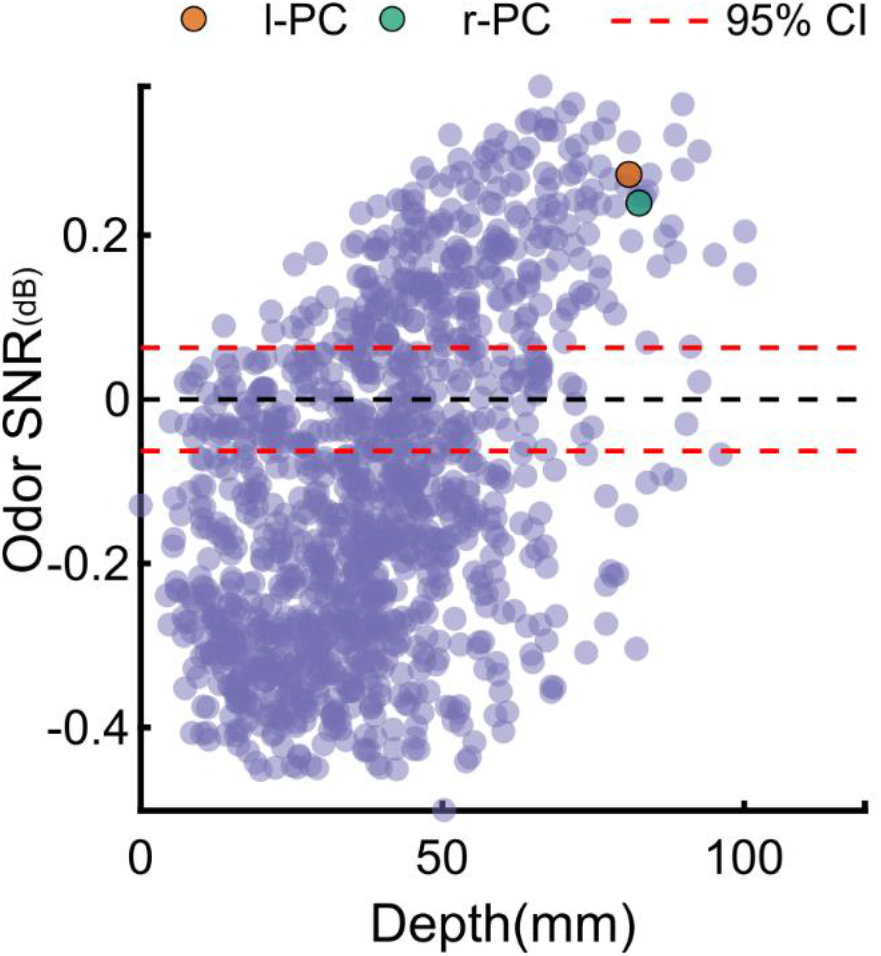
Signal level in the PC source reconstruction vs. other possible surrounding sources. Odor related SNR level as a function of depth with each point in graph showing the level of SNR for a possible dipole in the brain. The black dashed line marks 0, where the level of signal and noise are equal. Red dashed line shows 95% CI at the depth 80~100 mm which corresponds to PC depth. Left/right PC shown by orange and green circles, respectively, where both demonstrates SNR levels well above upper bond of noise CI.

### Source Connectivity

To provide a full picture of OB-PC connectivity during odor processing, we characterized the functional and effective connectivity between OB and PC using two separate, yet related, analyses. In the following section, we first explain the functional connectivity analysis in which the reconstructed OB and PC time-courses were transformed into the Fourier space. This data was subsequently used to assess the cross spectrogram as a measure of functional connectivity. This allowed us to identify frequency and time points where linear information transfer occurred between the OB and PC. Next, using spectrally resolved Granger causality, we assessed if the relationship of OB-PC was casual (i.e. the effective connectivity between OB and PC). This was done by transforming reconstructed signals to the Fourier domain and calculating a transfer function to estimate the spectrally resolved granger causality (Dhamala et al. 2008).

### Functional connectivity in frequency and time

Auto and cross spectral density of OB and PC were estimated by means of multi-tapered convolution method, implemented in the Field trip toolbox 2018 within Matlab R2019b (Oostenveld et al. 2011). Odor and Clean Air trials were separately transferred to Fourier space with 2 tapers from discrete prolate spheroidal sequences (DPSS) using a flexible time window that captures at least two cycles [20 ~ 1000ms] of each frequency bin. The frequency smoothing parameter was set to 80% of the targeted frequency (**Fig 2**). Given the inherent inability to achieve high sensitivity in both the temporal and frequency dimensions, this approach allowed us to have maximum time resolution of estimation but naturally smoothed the frequency dimension, proportional to frequency values, and thus more in gamma bands. However, because the gamma band is defined as a broad band (30~100 Hz) and odor processing in humans are not thought to occur in the higher range (Iravani et al. 2020), we prioritized high sensitivity in the time domain.

### Effective connectivity in frequency domain

The frequency content of the source time-course during the 1 second stimulus was estimated for [0~100 Hz] with a step of 1Hz using multi-tapered fast Fourier transform with the smoothing parameter of 4Hz and 7 tapers from DPSS. Contrary to the functional connectivity analysis, in this effective connectivity, the window length is equal to total length of stimulus interval (i.e. 0-1 s), therefore we could afford low smoothing parameter and consequently achieve high frequency resolution. The multivariate spectrally resolved granger causality measures were estimated at the individual level by computing the transfer function of OB to PC and PC to OB from cross spectral density using Wilson-Burg algorithms (**Fig 2**)(Dhamala et al. 2008). Subsequently, to increase the statically power, we averaged the two hemispheres. The statistical significance of effective OB-PC connectivity on the group-level was finally determined by two-tailed student t-test across subjects.

### Supported vector machine learning

To assess whether odor information is conveyed by the temporal dynamic of OB-PC connection, we used support vector machine learning (SVM) to classify odors (3 odors) from the information provided by the level of association between OB and PC on the group level. One individual did not respond in the identification task and was subsequently removed from the SVM analysis. Hence, data of 28 individuals was used in this analysis to create a subject level OB-PC connectivity maps for each odor. To construct the feature space from cross spectral density, the measure of OB-PC connectivity, a neighbor of 5 and 5 samples were considered for frequency and time dimension, respectively. Therefore, for each bin of the cross spectral density map neighboring datapoints in the time-frequency plane within the distance of 5 samples in each axis (i.e., time, frequency in both directions) were used to create the feature space. Given the distance of 5 samples in both directions, 11 time and 11 frequency bins were considered as the neighbors and used in the feature space. Hence, a total of 121 time-frequency pairs were used for each bin in the cross spectrogram for estimating the classification accuracy. The bins that had fewer than 10 neighbors were excluded from further analysis. Next, the whole cross spectrogram was assessed in a searchlight manner.

Features were unity normalized and the data was partitioned into one-leave-out scheme with all the three odor stimuli conditions counter balanced, spanning all the cases where each subject were left out at least once. Next, we classified the 3 Odor stimulus conditions. The mean accuracy on the group level was compared with the chance level (.33; given 3 odor categories) using non-parametric statistics, 5000-permutation Monte Carlo test. Finally, the accuracy for the individual odors were extracted from the time-frequency point found in the mean accuracy map. Given that each odor has a slightly different latency and frequency representation in human OB (Hughes et al. 1969), we allowed for ±40ms jitter in latencies and ± 10 Hz in frequencies for estimating the individual odor accuracies.

## RESULTS

### Early fast, and late slow, functional connectivity between olfactory bulb and piriform cortex

We first set out replicate our past finding, that the OB initially processes odors in the gamma frequency (Iravani et al. 2020), to assure that the EBG method extends also to this dataset. For this analysis, we used maximum frequency resolution to validate our OB source extraction method whereas subsequent analyses described below maximize temporal resolution. As previously published, the OB demonstrated initial oscillations in the gamma range (**Fig 4A**). Having replicated our past finding, we first assessed participants ability to correctly identify all odor presented to them during the experiment to assure that odors could be identified by name. We found that all participants but one (whose respond file was corrupted) were able to correctly identify the odors, and Clean Air as such, with high accuracy (mean: 89% ± 9%). We then assessed the functional connectivity, measured as information transfer in cross spectral analysis, between OB and PC during the odor presentation using cross spectrograms. The reconstructed OB and PC time-courses were transformed into Fourier domain and the auto and cross spectral density was estimated using multi-tapering convolution method comparing Odor to Clean Air conditions (inhalation of odorless air). In the OB, we found, as expected, initial gamma activity followed by beta activity (**Fig 4B**), whereas activity in lower frequencies (theta and beta) were found for the PC (**Fig 4C**). More importantly, assessing functional connectivity between OB-PC, we found a temporal transition across frequencies when assessing the cross spectrogram (**Fig 4D**). The earliest odor related functional connectivity was demonstrated around 100ms in high gamma ~70Hz, *t*(28) = 2.131*, p* < .042*, CI* = [0.003 0.151], which then evolved to slower oscillations around 500-700ms in low gamma ~35Hz, *t*(28) = 2.32, *p* < .028, *CI* = [0.013 0.200], to beta (~16 Hz) around 740-840ms, *t*(28) = 2.466*, p* < .020*, CI* = [0.018 0.194], and transferring to theta/delta (3 Hz) at later time points around 670-1000ms, *t*(28) = 2.620, *p* < .014, *CI*=[0.031 0.257](**Fig 4D**). There are multiple reports of laterality differences in odor processing (Royet and Plailly 2004). In a next step, we therefore separately analyzed processing in the left and right OB-PC connectivity. Here, we found a similar pattern for OB-r.PC and OB-l.PC. However, comparing Odor with Clean Air for OB-l.PC we find a weaker early gamma, *t*(28) = 1.27, *p* > .21, but significant late beta, *t*(28) = 2.32, *p* < .028 *CI* = [0.022 0.345] and theta/delta *t*(28) = 2.68, *p* < .012, *CI* = [0.032 0.241] (supplementary **Fig S2**). Further analysis indicated that the difference in early gamma is potentially driven by the Clean Air, *t*(28) = 2.03, *p* = .05 rather than Odor, *t*(28) = 1.54, *p* > .13 (supplementary **Fig S3**), although this potential effect was not significant.

**Fig 4.**
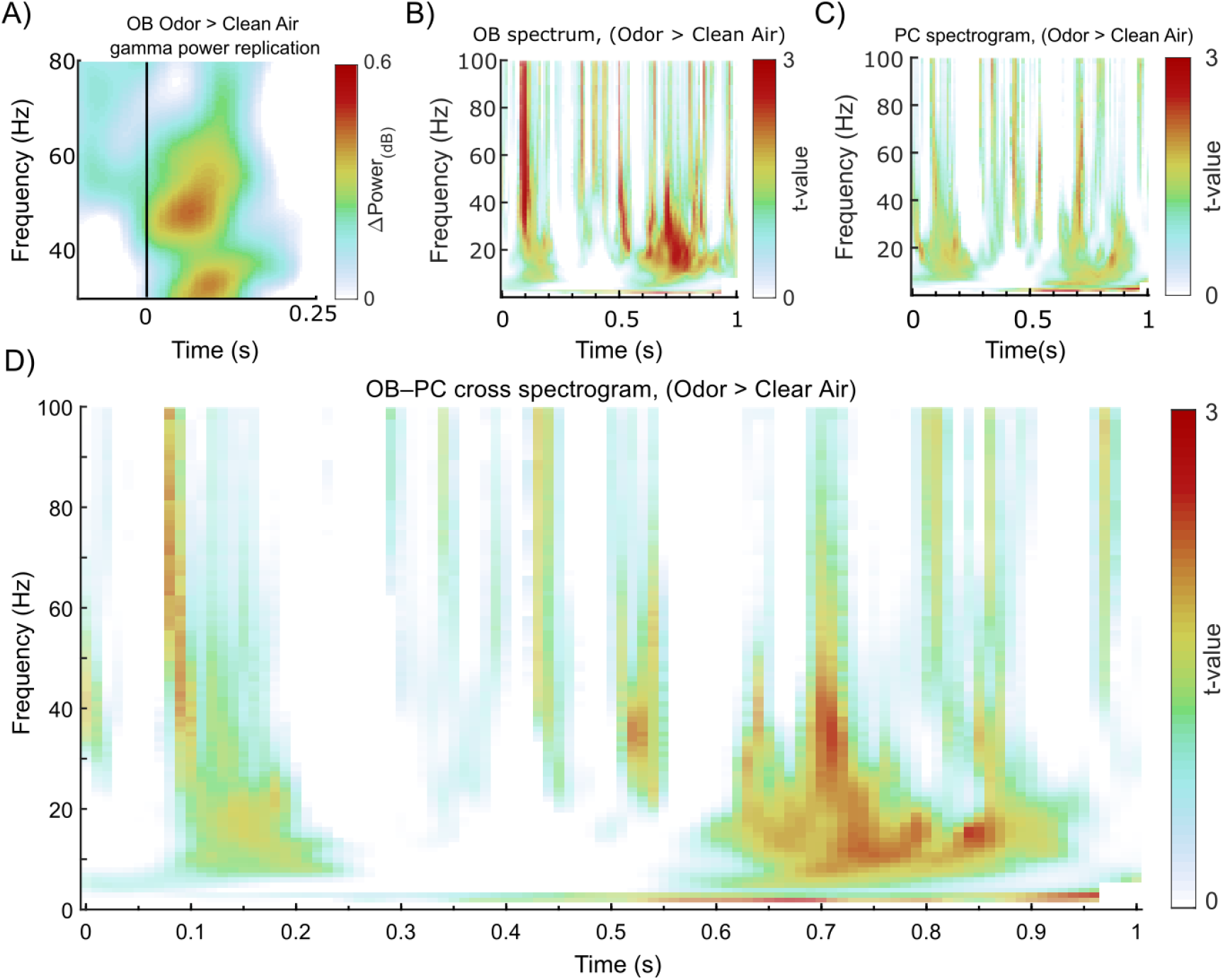
Auto and cross spectrogram of OB and PC. (**A**) Heatmap shows OB power spectrogram with finer frequency, rather than temporal resolution, to replicate original finding of OB processing (Iravani et al. 2020) (**B**) Heatmap of t-values for the spectral density of olfactory bulb (OB), when compared Odor against Clean Air. (**C**) Similarly, spectral density for Piriform cortex (PC) when comparing Odor against Clean Air. (**D**) The cross spectrogram shows frequency and time points where OB and PC are related, or functionally connected more for Odor compared with Clean Air. In panels b-d, t-values are color coded according to color scale on right side of figure using identical scale.

### Reciprocal effective functional connectivity between olfactory bulb and piriform cortex

Connectivity between two neural populations can be described either as functional or effective. Functional connectivity refers to the mere statistical dependency of amplitude (Kaboodvand et al. 2018; Biswal et al. 1995) or phase of signal between two populations (Kaboodvand et al. 2020; Kaboodvand et al. 2019) whereas effective connectivity refers to a predictive relationship between two populations (Eldawlatly and Oweiss 2010). To assess the effective connectivity between OB and PC, we used the frequency domain of Granger causality, a popular method for assessing if the future of a time series *x* can be predicted of the past of time series *y* over and above what can be predicted from the past of *x* alone (Granger 1969). This allowed us to characterize the function of olfactory circuitry in a directed manner both in the time and frequency domains (Seth et al. 2015). The reconstructed time-courses of bilateral OB and PC were transformed into the Fourier space by multi-tapered fast Fourier algorithm. In the frequency domain, the relationship between OB and PC was assessed as a function of frequency for both afferent versus efferent directionality (**Fig 5A, B**) and Odor versus Clean Air (**Fig 5C, D**) using multivariate spectrally resolved Granger causality. Benefiting from the directionality of the Granger causality method, we found that higher frequencies with peaks in the beta range around ~30Hz, *t*(28) = 2.953*, p* < .006*, CI* = [0.208 1.150], and gamma around ~58 Hz, *t*(28)=2.865*, p* < .008*, CI* = [0.148 0.888], facilitated the afferent connection (i.e. from OB to PC; **Fig 5A**). For the reverse connection, the efferent link from PC to OB, we only found connection in the lower delta frequency, *t*(28) = 5.074*, p* < .0001, *CI* = [1.076 2.533], and theta frequencies around 6Hz, *t*(28)=2.078*, p* < .047*, CI* = [0.011 1.605], during odor processing (**Fig 5B**). There was no significant relationship in signals from OB to PC induced by a inhalation (Clean Air > Odor) except from a connection in a narrow band around 60-70 Hz, *t*(28) = 2.145, *p* < .041, *CI* = [0.019 0.809] (**Fig 5C**). Likewise, no significant inhalation-related relationship was found for the reverse direction, from PC to OB (**Fig 5D**). There was no absolute effect of laterality for the effective connectivity, neither from OB to PC or vice versa, or for neither the Odor nor Clean Air-Odor contrast, except for a single peak at ~55 Hz, *t*(28) = 2.74, *p* < .011, *CI* = [0.08 0.59], that was stronger for right rather than left hemisphere when Odor was contrasted against Clean Air (supplementary **Fig S4**).

**Fig 5.**
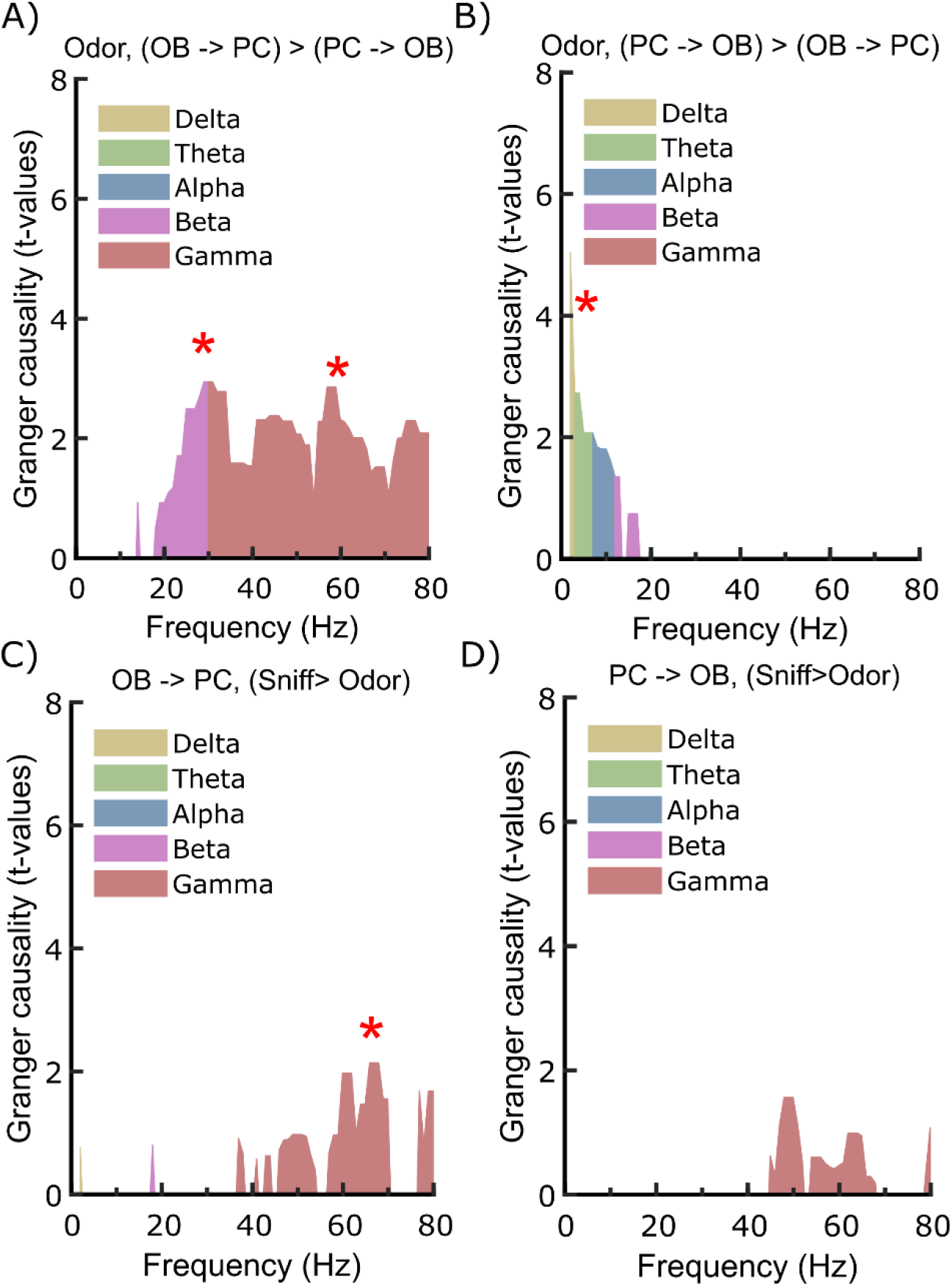
Effective connectivity between OB and PC for incoming versus outgoing during Odor and Clean Air. (**A**) Spectrally resolved Granger causality analyses during the 1 second odor presentation show significant effective connectivity from OB to PC in the beta and low gamma band for Odor compared to Clean Air. (**B**) Significant connectivity is found from PC to OB in slower delta/theta band for Odor compared with Clean Air. (**C**) Connectivity from afferent, OB to PC, shows a significant increase in effective connectivity for Clean Air vs Odor in a narrow band around 60-70 Hz. (**D**) No relationship found for Clean Air from efferent, PC to OB. In all panels, colors indicate frequency band division and red star in graph indicate significant peaks.

### Connectivity between the olfactory bulb and piriform cortex in low gamma reflects odor identity

We found reciprocal causal relationships between OB and PC, but in different frequency bands. The afferent connection from OB to PC was found to be in broad band beta/gamma and efferent link, from PC to OB, in slow delta and theta band. To assess whether patterns of this connectivity could be linked to odor, we further tested if our included odors could be read out from OB-PC connectivity using support vector machine (SVM), a supervised learning approach. The main prediction was that if content-specific representations are contained within the OB-PC connectivity, then the SVM classifier should significantly differentiate between the three odors. Alternatively, if the OB-PC connectivity is due to a non-specific effect of odor stimulation, then the classifier should not be able to differentiate between the odors. In addition, we were interested not only if, but also when, information about the odor might be transferred. Therefore, we used SVM on the cross spectrogram of OB-PC to identify frequency/time points where the odor identify can be read out above the chance level. To facilitate this approach, we unity-normalized values from the cross spectrogram for each condition and assessed each map during 1 second stimulus and broad band (1~100 Hz) in a searchlight framework on the individual level. When assessing the cross spectral density of OB-PC using SVM, we found that a time-frequency window around 100ms and 35-45 Hz allowed us to dissociate the three odors. Mean classifying accuracy within this window was significantly above chance level with peak of mean accuracy being .42 (**Fig 6A**), *t*(27) = 3.29, *p* < .002, and probability confidence interval range *CI-range* = 0.001 using 5000 Monte Carlo permutations test (**Fig 6B**). In addition, we found that an extended area around 300ms after odor onset in the 50-70Hz frequency range, a small area at the same time around 30Hz, and intermitting time-periods in the theta band facilitated classification. The slight differences between the descriptive map (**Fig 6A**) and statistical map (**Fig 6B**) might possibly be due to non-normal distribution of accuracies that has been resolved by the non-parametric Monte Carlo permutations test in this analysis. To further assess whether this classification was due to spurious non-specific effects, we repeated the classification but for connectivity between the OB and postcentral gyrus (PCG). We selected PCG because it is an area that demonstrate low functional connectivity probability with piriform cortex in the large online Neurosynth database (www.neurosynth.org) yet has been demonstrated to process non-odorous intranasal stimuli. We found no above chance classification accuracy around 100ms and 35-45Hz, thus suggesting that the odor classification within this time window is specific to OB-PC connectivity (**Fig 6D, 6E**).

**Fig 6.**
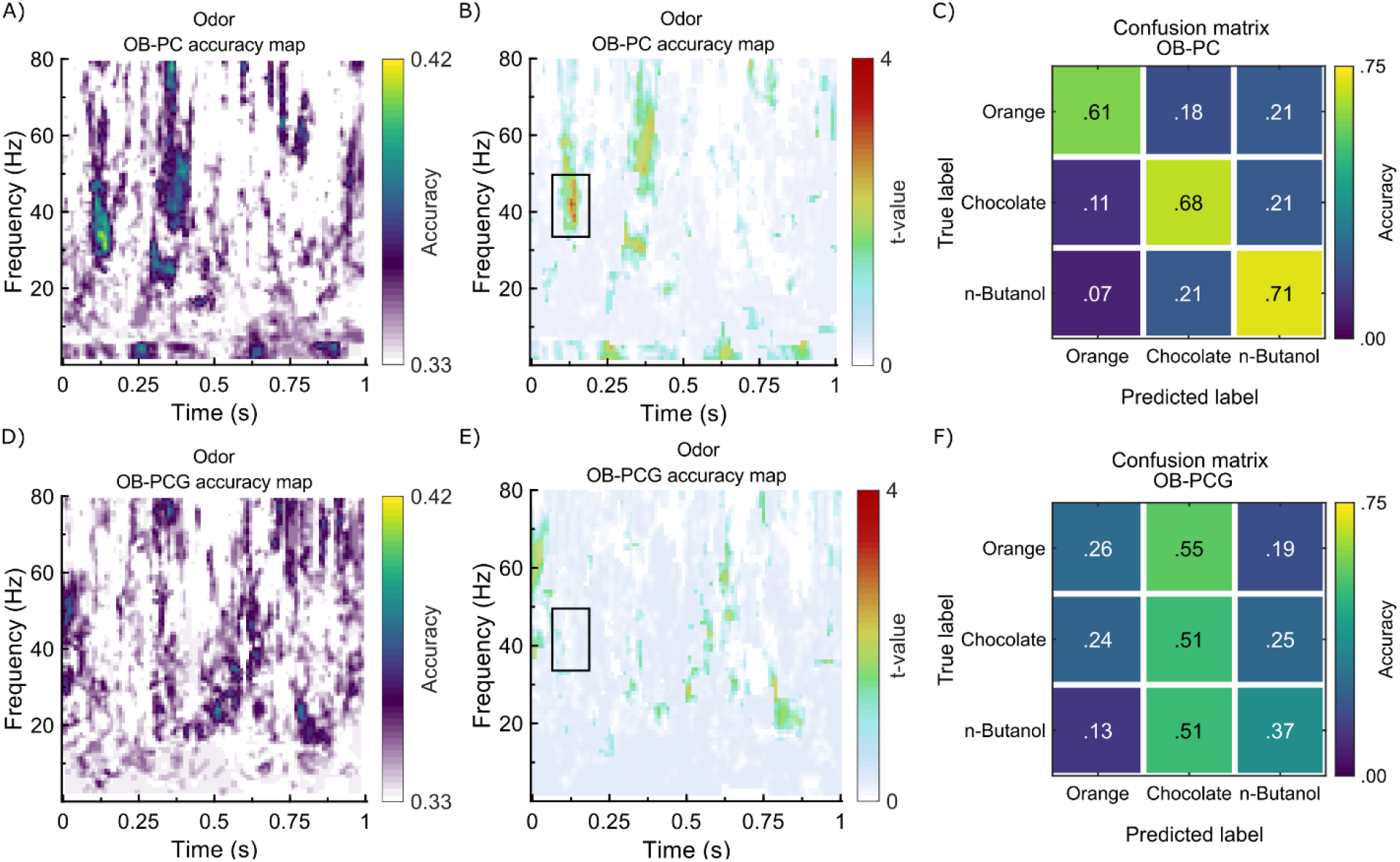
Odor identity is read out from OB-PC connectivity. (**A**) Heatmap shows the accuracy for classifying 3 odorants by assessing the <u>connectivity of OB-PC</u> using SVM. (**B**) Heatmap shows the t-map of accuracy for the connectivity of OB-PC. The accuracy of 4 standard deviations above chance level (.33) was achieved around 100ms post odor onset and 35-45 Hz shown by black box. (**C**) The confusion matrix for the OB-PC connectivity shows the accuracy for each individual odor. n-Butanol, Chocolate, and Orange were classified with accuracy .71, .68 and .61, respectively (**D**) Heatmap shows the accuracy for classifying 3 odorants by assessing the <u>connectivity of OB-PCG</u> using SVM. (**E**) The t-map accuracy is shown as the heatmap, where we found no accuracy above chance level within the time-frequency of interest shown by black box where the odor could be read out from OB-PC connectivity. (**F**) The confusion matrix for the connectivity of OB-PCG show an indiscriminative arrangement.

We further assessed the accuracies for the individual odors in the area where accuracy was above chance in the OB-PC connectivity map. We found .71, .68 and .61 accuracy for n-Butanol, Chocolate, and Orange, respectively (**Fig 6C**). Repeating the same analysis for OB-PCG connectivity map showed indiscriminative patterns for odors (**Fig 6F**).

## DISCUSSION

Here we have captured the functional connectivity of the human OB and PC during odor perception. We show that this dynamic connectivity is based on a wide range of oscillatory frequencies that change as a function of the direction of the signal and the time from odor onset. We demonstrated that odor can be decoded from the OB-PC oscillatory connectivity as early as 100ms after odor onset, thereby suggesting that odor identity, here defined as the identity of the odorant rather than its associated object, is at least in part deciphered in OB-PC oscillatory communication. Importantly, these findings show that while human and non-human animals share commonalities in OB-PC connectivity, the human OB-PC connectivity has partly different frequency distribution.

Source reconstruction from human surface EEG is spatially less specific than, for example, surgically placed intracranial electrodes; although, EEG as a method was recently demonstrated to be a reliable method for extracting radial and deep sources (Piastra et al. 2021). That said, we have previously shown and extensively validated in several experiments that the EBG signal can be effectively extracted and that it reliably originates from the OB (Iravani et al. 2020). Indeed, here we replicated our past finding of an early (<150ms) odor-dependent OB signal in the gamma band. The PC dipole is more difficult to validate. Our sensitivity analyses did, however, demonstrate that our PC sources achieve higher odor-dependent SNR than other possible source-locations in the surrounding area. Moreover, in our analyses of the PC spectrogram in response to odor stimulation, we partially replicated previous findings obtained from intracranial electrodes implanted in the human PC (Jiang et al. 2017). Using these two dipoles, we found reciprocal communication between OB and PC in different frequency bands. Except weak gamma afferent communication, our result indicates a lack of communication between OB and PC during the no odor trials which emphasize the dependency on the casual communication between OB and PC in response to odors. Moreover, afferent connections from OB to PC during odor stimuli seemed to utilize mainly the gamma and beta band whereas the efferent, top-down connection operated primarily in the theta band. Differences in frequencies for afferent and efferent connections have previously been reported in both the olfactory (Fourcaud-Trocmé et al. 2019; Kay 2014) and visual systems (Bastos et al. 2015; van Kerkoerle et al. 2014).

Generally, in the animal model literature, higher frequencies, such as gamma, has been linked to within area processing and afferent ‘bottom-up’ communication whereas lower frequencies, such as beta, have been linked to efferent ‘top-down’ communication (Richter et al. 2017; Bastos et al. 2015; Frederick et al. 2016). Indeed, odor stimulation produces OB-PC coherence in the beta band, which is mainly induced by efferent communication (Neville and Haberly 2003; Gray and Skinner 1988; Martin et al. 2006). However, others have found afferent communication from the hippocampus to the OB in the beta and theta band (Gourévitch et al. 2010) where the beta communication is more relevant for before odor sampling (Kay and Freeman 1998), suggesting that it is a too simplistic notion that beta is restricted to efferent communication. Whether these discrepancies are due to differences in task-demand (Beshel et al. 2007; Frederick et al. 2016) or region-specific effects is currently unclear but from a biophysics point of view, the slower beta oscillations are more suited for long range transmission of information between areas, such as between OB and hippocampus (Kopell et al. 2000). Likewise, the use of non-invasive recording methods might promote slower oscillations and it worth noting that these signals are obtained in healthy humans. Nonetheless, we demonstrated here that in humans, afferent communication from the OB to PC is dominated by the gamma band but also activity in the high-beta band. Although it is unclear whether it constitutes a meaningful difference, it is interesting to note that this beta band activity is in the higher range (around 30Hz) of what is commonly observed in animal models where 15-30 Hz are commonly reported in the literature (cf. Martin and Ravel 2014). It is not clear, however, what impact the behavioral tasks used in the different studies might have for the demonstrated differences in odor processing between the human and non-human animal literature.

One possible reason for our finding of beta oscillatory involvement in both afferent and efferent communication in humans is the difference in respiration pace between humans and rodents. The respiratory and olfactory systems are linked and respiratory cycles in humans are significantly slower compared to rodents (Mainland and Sobel 2006; Rojas-Líbano et al. 2014). Moreover, an intercranial study in humans found that the inspiratory cycle entrains oscillatory activity in PC, demonstrating that respiration, and potentially the respiration pace, moderates the neural activity in PC (Zelano et al. 2016). Given the tight coupling between breathing phase and odor perception, where orthonasal odors are primarily experienced during the inhalation phase, this difference may result in species-specific differences in communication frequencies within the olfactory system. In line with this notion is a recent evidence that cross-frequency coupling is dependent on respiration pace (Hammer et al. 2020). A further indication supporting this notion is our results demonstrating that efferent communication from the PC to OB during odor stimulation was dominated by theta oscillations, a frequency that in the animal literature has been linked to respiration (Kay 2014). Interestingly, a recent intracranial study in human participants undergoing elective surgery for intractable epilepsy provided further evidence for odor related theta activities by demonstrating that odor processing within the PC is dominated by theta activity (Jiang et al. 2017). Together, this shows that to fully understand similarities and differences in processing, translational studies comparing model organisms with human participants using identical tasks and odors are needed.

Past studies have demonstrated that odor can be decoded from activity within the PC assessing both patterns of distributed piriform neurons in rodents (Roland et al. 2017; Rennaker et al. 2007) and oscillatory signals (Jiang et al. 2017) as well as summated neural responses in humans (Howard et al. 2009). Here we demonstrated that it is also possible to decode odor from the OB-PC connectivity as early as 100ms after odor onset. These types of analyses lack directionality but given the early time of the identified decoding cluster and our granger causality analyses demonstrating bottom up directionality in this frequency band, it is likely that our analyses are tapping into information from the OB to the PC. However, it should be noted that even though we identified a homogeneous and extended cluster in the 35-45Hz frequency, we only had 42% classification accuracy. Although this is significantly different from chance level (33%), it is not a strong result; potentially due to the low number of odors, thereby the confinement of available perceptual space was not allowing a clear odor differentiation. Using a larger battery of odors in future studies might provide a better classifying accuracy. Alternatively, our results might link to the unique perceptual experience of these three odors and not generalizable to other odors. Nonetheless, the latency of the decoding results corresponds with past intracranial recording from the PC where odor related activity was found within 500ms (Jiang et al. 2017) and also occurs at a time point that is close to the initial OB processing response (about 100ms past odor onset) that we have shown earlier using a similar method (Iravani et al. 2020), and replicate in this data (**Fig 4a**). However, the above chance performance extends to two temporal windows around 300ms for gamma band and reverberate during the full 1s for theta band (**Fig 6B**). Although speculative, these findings fit well with the directed Granger connectivity results (**Fig 5A, 5B**). Additionally, our decoding finding is further strengthened by the fact that we were not able to decode odor from OB connectivity within the same time period based on connectivity with the control region (i.e., PCG), thus supporting the specificity of our OB-PC finding. However, when we separately assessed the accuracy for odors, we obtained the maximum accuracy of 71% for n-Butanol whereas the accuracy for Chocolate and Orange were found to be 68% and 61%, respectively, suggesting heterogeneity in classifier performance across odors; perhaps due to differences in the latency and frequency of odor representations in the OB-PC connectivity. Moreover, the non-specificity and high dimensionality of the employed classification scheme unfortunately makes it difficult to deduce exactly what aspect of the odor that is coded in the OB-PC connection. When we assessed the perceptual aspect of odor, we only found the effect for pleasantness. However, this differences in the pleasantness did not modulate the inhalation parameters, length of inhalation or area under the curve. Given that inhalation response is such a robust measure of odor valence that is used as clinical test (Frank et al. 2003), the finding of no difference in the inhalation parameters as function of odors suggest that potential effects of pleasantness cannot have a decisive influence on the results. We argue that the surprisingly large difference in pleasantness ratings between odors might originate from a contrast effect given that our odor stimuli did not span a large section of available perceptual space. Additionally, assessing the confusion matrix brought to view that two pleasant odors are not confused by the decoding method, hence, making it unlikely that pleasantness is an underlying parameter that is encoded in the OB-PC connection. The question regarding what exactly is decoded might be better assessed in animal models where fewer trials are needed to obtain high SNR, or in human experiments with a large and diverse set of odors that vary in molecule structure, quality, associations, and valence and therefore the contribution of each factor can be systematically explored.

In summary, using a novel, non-invasive technique for assessing activity in the first central relay in the olfactory pathway, we demonstrated here that the olfactory bulb and piriform cortex, two essential nodes in the human olfactory system, show reciprocal, oscillation-based, communication in a frequency- and time-dependent manner during odor stimulation. In response to an odor stimulus, the olfactory bulb’s afferent communication to the piriform cortex is dominated by oscillation in the gamma and beta bands. Top-down piriform cortex input to the olfactory bulb, in contrast, was dominated by theta band oscillations. Moreover, we demonstrated that odor identity could be decoded from this reciprocal interaction within 100ms of odor onset. These data further our understanding of the critical role of bidirectional information flow in human sensory systems to produce perception.

## ACKNOWLEDGMENTS

Funding provided by the National Institute on Deafness and Other Communication Disorders (R21DC016735) as well as the Knut and Alice Wallenberg Foundation (KAW 2018.0152), awarded to JNL. AA is supported by a grant from the Swedish Research Council (2018-01603).

## DATA AND CODE AVAILABILITY

We collected EEG/EBG data from 29 healthy individuals. We publicly share all data and code for the main analyses of the data used in this study at an open science framework. Data and codes are available for download on https://osf.io/j7fae/?view_only=7194144d5c7d4ddc9fd33ce7e483afa5

## Footnotes

1 We are here using the term identity to indicate the identity of the odorant or mixture rather than linking the term to an ability to name or otherwise process the odor object the odorant or mixture has become associated with.

