## supplementary Fig for "ODOR IDENTITY CAN BE EXTRACTED FROM THE RECIPROCAL CONNECTIVITY BETWEEN OLFACTORY BULB AND PIRIFORM CORTEX IN HUMANS"

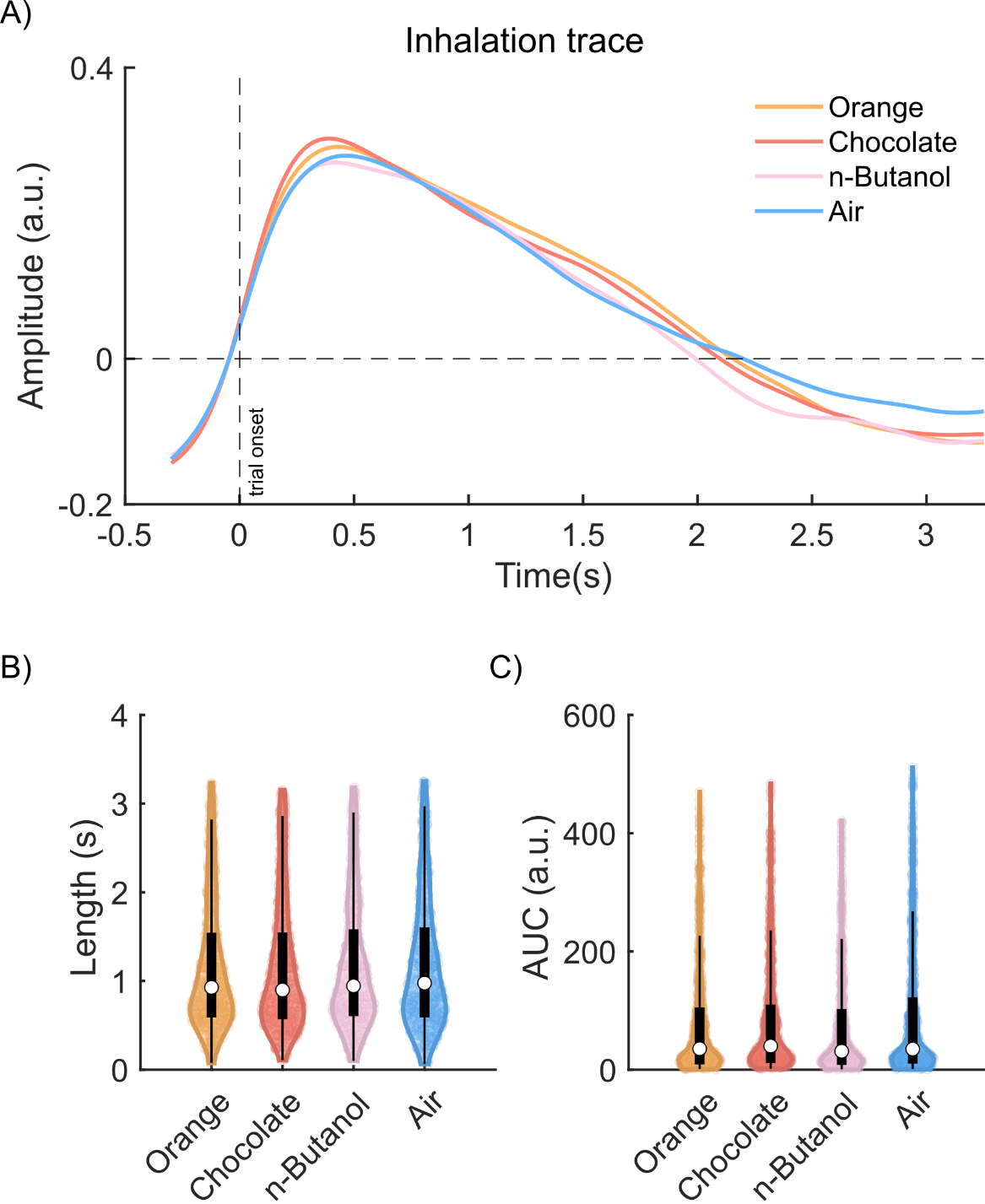

***Fig S1. Breathing parameters.*** *(****A****) Averaged inhalation trace for all four conditions showed similar pattern. (****B****) The violin plot shows the distribution of inhalation’s length for different conditions (****C****) Similarly to B violin plot shows the distribution of AUC of inhalation for different conditions. For more details on violin plots, please see the legend of Fig 1.*

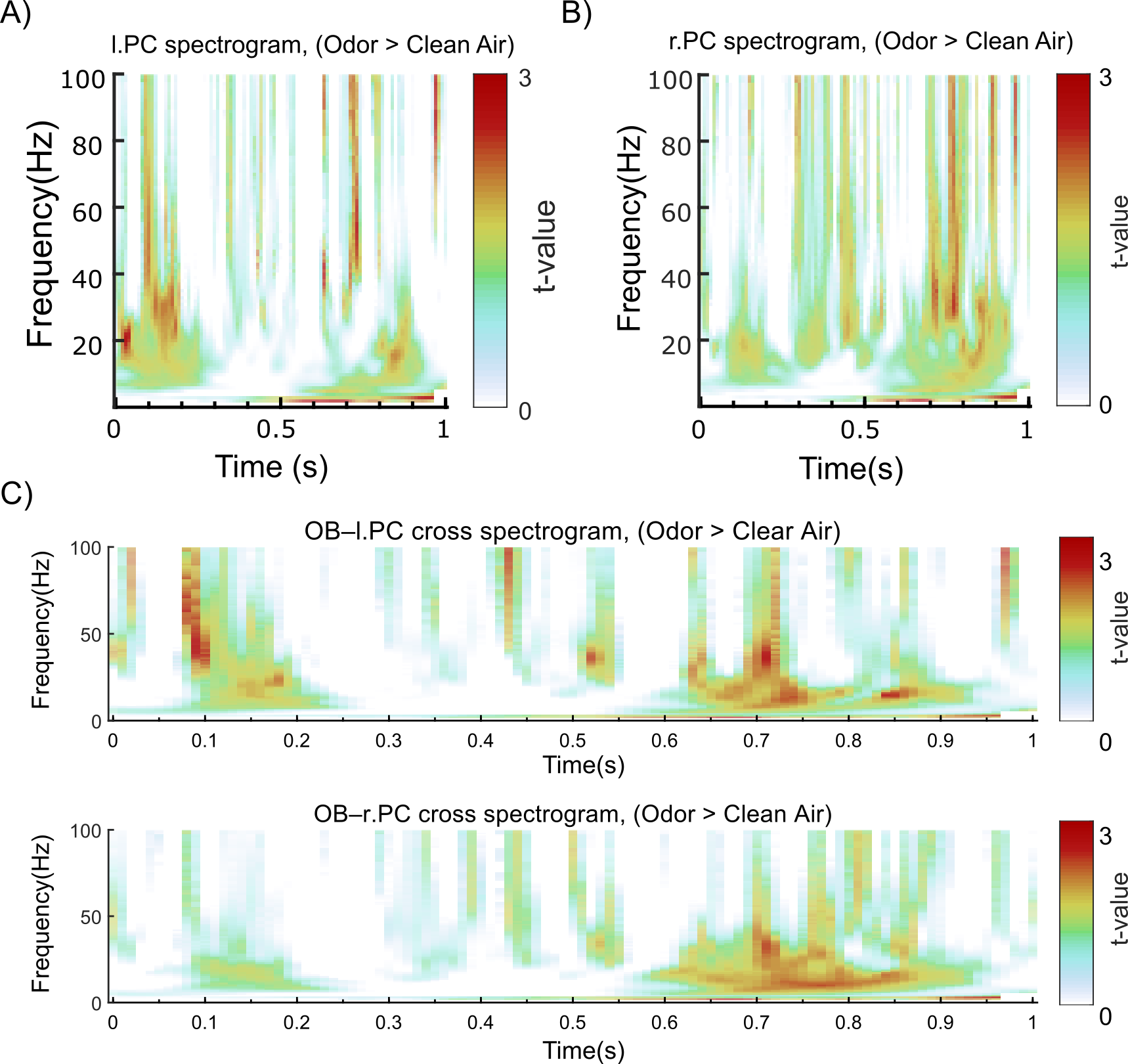

***Fig S2. Auto and cross spectrogram of left/right PC and averaged OB.*** *(****A****) Heatmap shows t-values for spectral density of left piriform cortex (l.PC) and, (****B****) for right piriform cortex (r.PC) when comparing Odor against Clean Air. (****C****) The cross spectrogram shows frequency and time points where OB and l.PC (upper panel) as well as OB and r.PC (lower panel) are functionally connected for Odor compared with Clean Air. In all panels, t-values are color coded according to color scale on right side of figure using identical scales.*

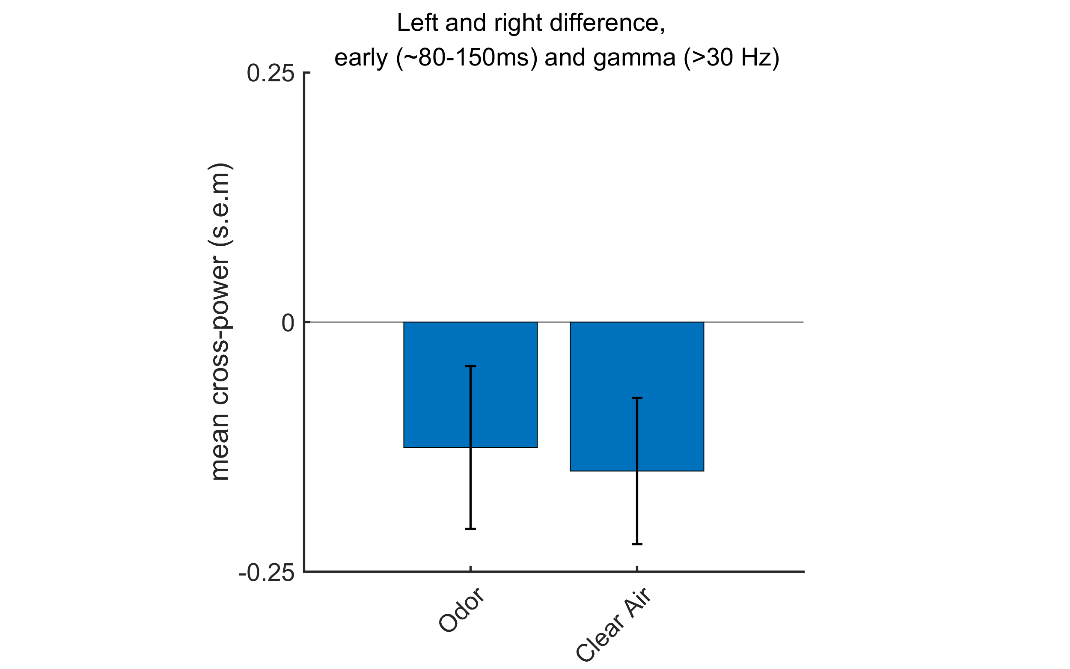

***Fig S3. Laterality effect of early gamma.*** *The bar graph shows the mean difference between left and right PC and OB connectivity during early gamma. No effect was found for Odor, t(28) = 1.54, p > .13, and only trends, t(28) = 2.03, p = .05, for Clean Air.*

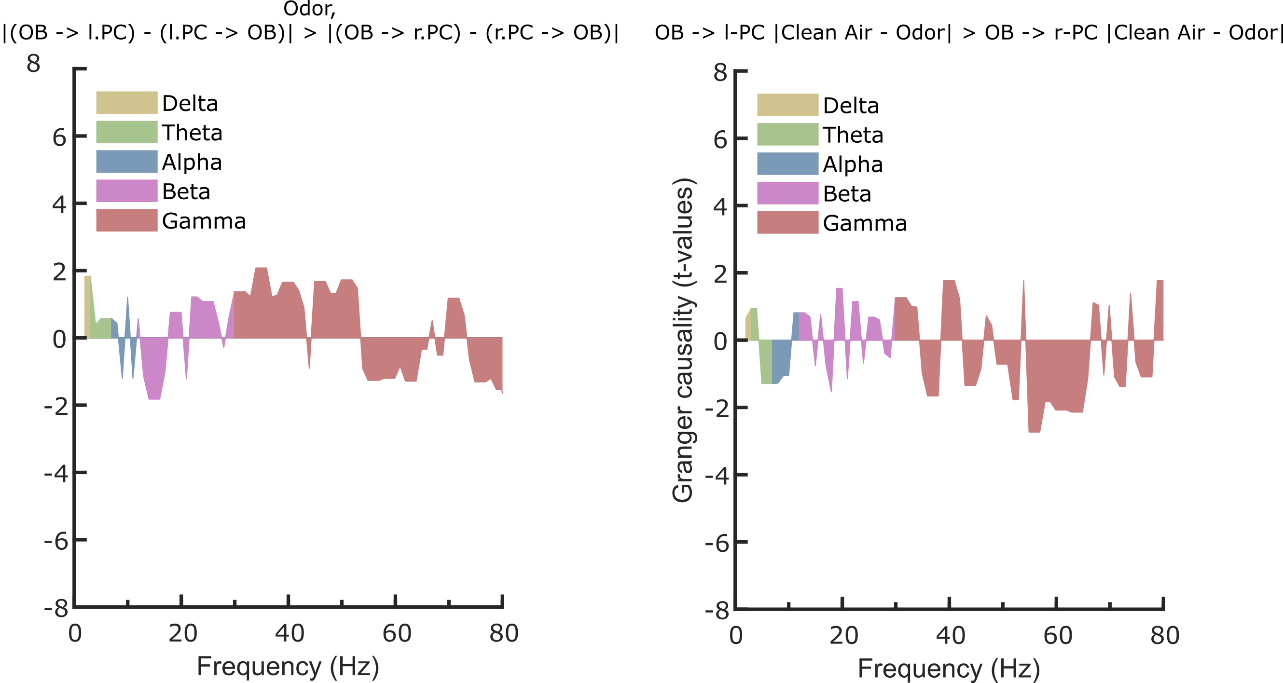

***Fig S4. Laterality and effective connectivity of OB-PC.*** *The spectrally resolved granger causality indicated no significant effect of laterality for odor trials. Likewise, the effective connectivity from OB to l.PC is not different from OB to r.PC for Odor and Clean Air contrast.*
